# A new tool for 3D segmentation of computed tomography data: *Drishti Paint* and its applications

**DOI:** 10.1101/2020.04.09.997965

**Authors:** Yuzhi Hu, Ajay Limaye, Jing Lu

## Abstract

Computational tomography is more and more widely used in many fields for its non-destructive and high-resolution in detecting internal structures of the samples. 3D segmentation of computed tomography data, which sheds light into internal features of target objects, is increasingly gaining in importance. However, how to efficiently and precisely reconstruct computed tomography data and better represent the data remains a hassle. Here, using a set of scan data of a fossil fish as a case study, we present a new release of open-source volume exploration, rendering, and 3D segmentation software, *Drishti* v2.6.6, and its protocol for performing 3D segmentation and other advanced applications. We provide new toolsets and workflow to segment computed tomography data thus benefit the scientific community with more accurate and precise digital reconstruction, 3D modelling and 3D printing results. Our procedure is widely applicable not only in palaeontology, but also in biological, medical, and industrial researches, and can be used as a framework to segment computed tomography and other forms of volumetric data from any research field.

## 1 Introduction

Nowadays, computed tomography (CT) is widely used in many fields, such as medicine, science, industry and education [1–5]. To have a better understanding of the data generated from CT, 3D scientific visualization becomes more and more important for researchers to obtain better insights [6–9]. For the last four decades, 3D scientific visualization field have been developed with new ways to visualize and analyse data more accurately [10–19]. Two techniques have emerged to provide different visualizations: surface rendering, the method of interpreting data-sets by generating a set of polygons that represent the wanted feature; and volume rendering [10], a more direct way for the reconstruction of 3D structures, which represents 3D objects as a collection of cube-like building blocks called voxels, or volume elements. However, the importance of scientific visualization software along with their functionality was underestimated by the community due to lack of public exposure and communications between multidiscipline.

3D segmentation, segmenting the internal structures in sequences of images, is a key tool for investigating and understanding the internal structures of target objects [6, 20–22]. The 3D segmenting methods rely on thresholding, edge detection, clustering, or region growing to group pixels based on brightness, colour, or texture [21]. Compared with researches carried out in the image processing field, more requirements have to be met towards building and developing applications which require the use of more efficient and effective techniques to save time and resources.

*Drishti*, as an open-source volume exploration, rendering and 3D segmentation software [23], is well-known for its superb visualization outcomes [6, 24–26]. Here we present the latest released *Drishti* v2.6.6, and the last developed module *Drishti Paint* v2.6.6, as a new toolset for 3D segmentation (2D and 3D painters) of volumetric data (i.e. a 3D volume produced by a group of 2D images of a slice or section of the scanned object acquired by X-ray imaging and Magnetic resonance imaging, etc.) by using the CT scan data of a fossil fish as case study. We also suggest protocols for how to perform 3D segmentation using *Drishti Paint* v2.6.6 efficiently and precisely. New features in *Drishti*, such as mesh generation and simplification, allow 3D printing, model simulation, etc.) are also covered in this study.

## 2 Materials and Methods

### 2.1 Materials

The material used in this study, a Devonian unnamed placoderm fish (ANU V244) from Burrinjuck, near Canberra, south-eastern Australia, is held at Australian National University [3], Canberra, Australia. The whole specimen and the separated right anterior upper toothplate were scanned in 2015 [27, 28] at CT Lab, ANU [7] using a HeliScan MicroCT system with 1.2mm aluminium/ 0.35mm stainless steel for the whole specimen and 2mm aluminium filters for the upper toothplate to yield sharp images [18, 29] at a resolution of 21μm and 2μm respectively. CT data was reconstructed using an in-house software called *Mango* (https://physics.anu.edu.au/appmaths/capabilities/mango.php).

### 2.2 Methods

The CT datasets were explored and rendered using *Drishti* v2.6.6. The individual data sets were read at low resolution, and contrast was incremented with the help of histograms and slides were filtered. The information thus generated is written as *.pvl.nc for saved files (i.e. processed volume format) or *. xml (i.e. Extensible Markup Language format) for saved projects. The scan data was demonstrated by a protocol for segmenting the 3D volumetric data using *Drishti Paint* v2.6.6. The right check complex was segmented out of the whole dataset. The internal structure under the region-of-interest of its right anterior upper toothplate has been segmented as another case study.

## 3 Results and discussions

3D segmentation is often hampered by the time-intensive nature of extracting information from samples, the complexity of preparing fossils digitally, and the need for hardware. Here we highlight the workflow and key features of the *Paint* module for *Drishti* and the general applications of *Drishti* using CT data of an Early Devonian placoderm fish [27, 28] as an example. A suggested workflow of segmenting volumetric data in *Drishti* v2.6.6 is presented here (figure 1), which can generate good results in terms of segmentation, rendering and surface mesh generation outputs. Our protocols can be used as a baseline or starting point to practice accordingly in order to obtain an ideal result for a specified dataset. To help with a better understanding into other aspects of *Drishti*, we also provide additional information about *Drishti*, such as installation instructions, a summary of all *Drishti*-supported import formats, additional learning resources and background of the program (see supplemental information).

**Figure 1.**
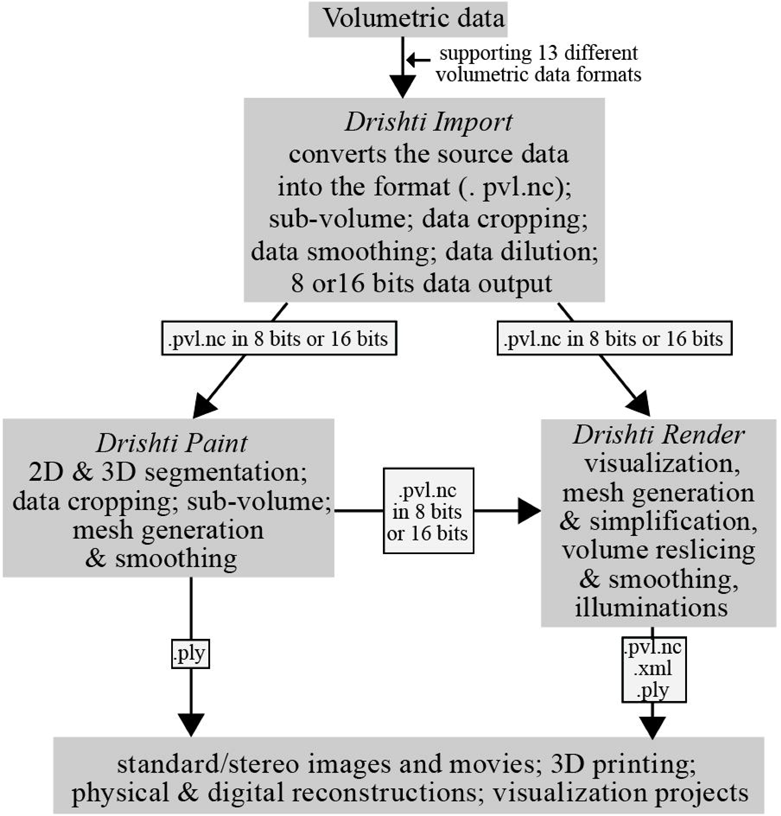
The general workflow for operating *Drishti* v2.6.6. Data formats in this figure are only abbreviations, for detailed information please refer to Supplementary Table 1.

Three modules, *Drishti import, Drishti render* and *Drishti paint*, have their capacities and features, and can be used in combination with each other to ensure an accurate and precise segmentation along with volume rendering to help with visualize a region-of-interest and solve scientific problems. *Drishti Paint* uses a combination of discontinuity detection based and similarity detection-based image segmentation approaches. There are two modes in *Drishti Paint* v2.6.6, Graph Cut and Curve, which are designed specifically to consider 2D slices in volumetric data or 3D volume directly.

Two transformations in mathematical morphology (i.e. erosion and dilation) were implemented in *Drishti Paint* and we used a combination of these two transformations with 3D Freeform Painter to help with a faster segmentation process. The right cheek complex has been segmented from the original CT dataset of the whole specimen (figure 2). The internal canal networks of a selected region-of-interest on the right anterior upper toothplate have been segmented from the original dataset of the toothplate (figure 3). Processed data then exported as separate volumes by extracting the tagged volume after tagging the regions of interests using the tagging function. Both segmentations were carried out in 16-bits full resolution in alignment with their source data and used type 2 gradient thresholding with values thresholding to select the range of the histogram for segmenting.

**Figure 2.**
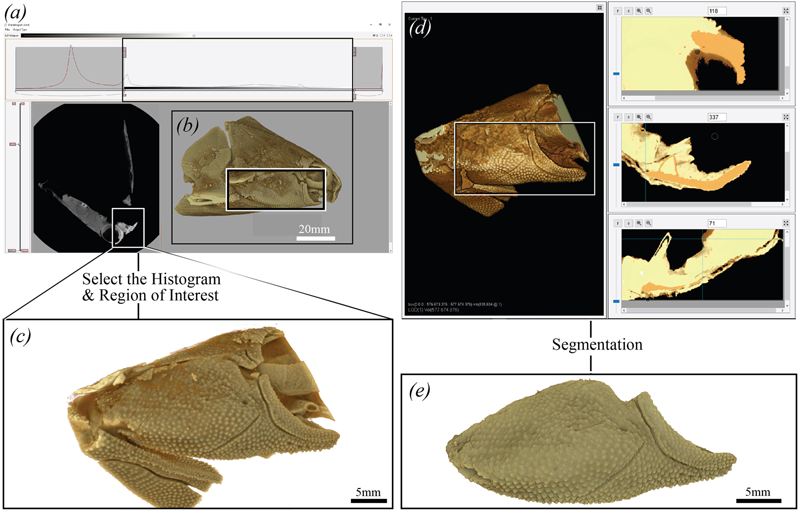
Extract a 3D virtual volumetric model in *Drishti*. A. Raw data of the right cheek complex of a Devonian placoderm fish (ANU V244) in *Drishti Import* v2.6.6. B. V244 in *Drishti Render* v2.6.6 in 16 bits full resolution (lateral view). C. Selected region-of-interest for segmentation. D. Selected region-of-interest in *Drishti Paint* v2.6.6 for segment the right cheek complex. E. Segmented right cheek complex in *Drishti Render* v2.6.6 (external view).

**Figure 3.**
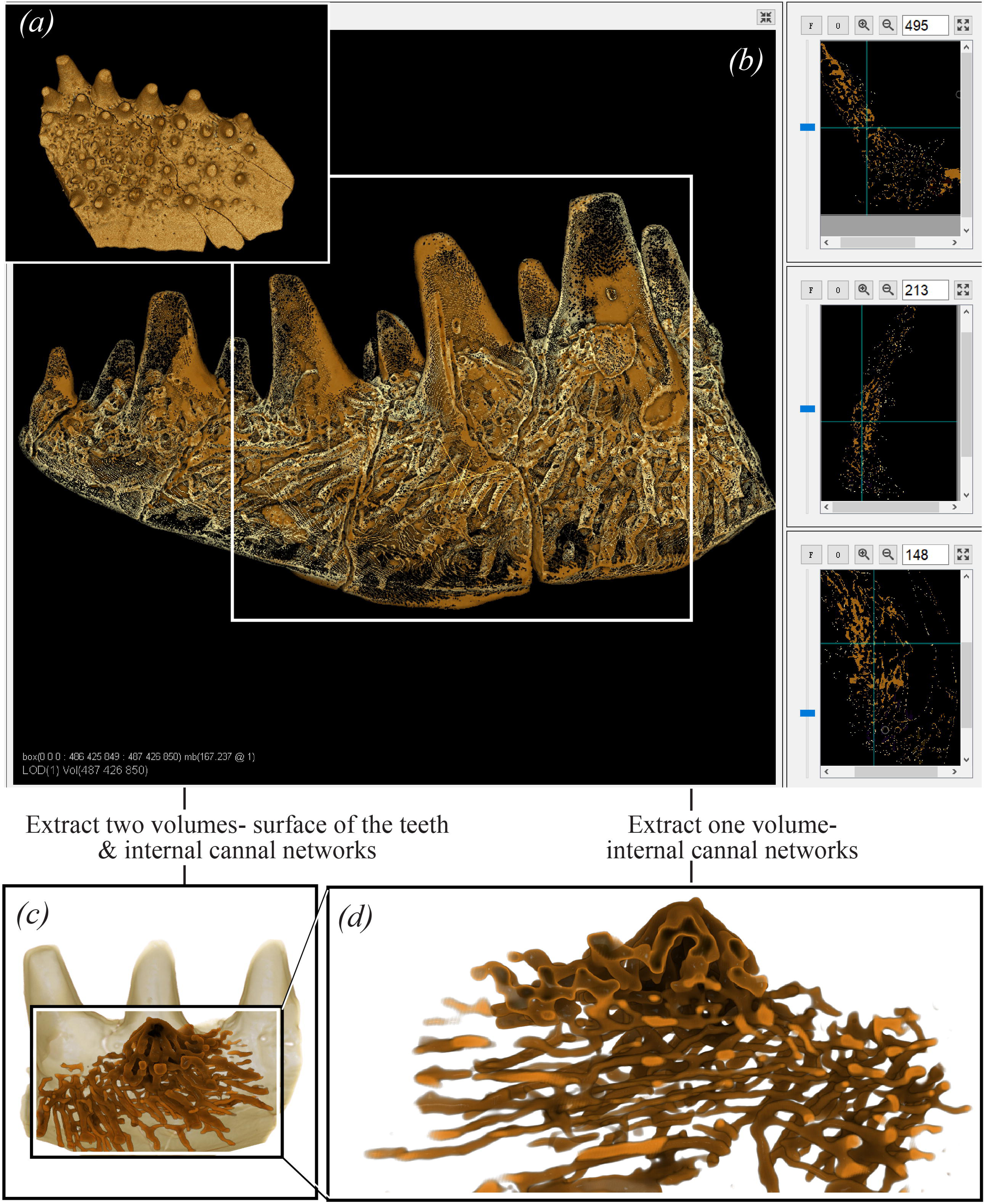
Segmenting internal structures of a toothplate (an Early Devonian placoderm fish, ANU V244) in *Drishti Paint*. A. Right upper anterior toothplate in *Drishti Render* v2.6.6 (external view). B. Raw data of the right upper anterior toothplate in *Drishti Paint* v2.6.6. The white-boxed area shows the selected region-of-interest on the right upper anterior toothplate for segmentation. C. Segmented outer surface of the region-of-interest on the right upper anterior toothplate and its internal canal networks underneath in *Drishti Render* v2.6.6. D. Segmented internal canal networks with a close-focus in *Drishti Render* v2.6.6.

We suggest that these two transforms are more useful when using clean and high contract dataset as a risk of changing the morphology of the input datasets by doing multiple transformations at once. Multiple-thresholding (i.e. use both values and gradient thresholding) in *Drishti Paint* v2.6.6 is very effective for volume segmentation and usually is the first step towards segmenting a volume (figures 2d and 3b), however, it is also very sensible to noise and intensity in homogeneities. In this case, to segment the internal canal networks of the selected region is tightly related to the selected thresholds, and any small changes in the thresholds values can lead to different segmented results, which is one of the key factors need to set up specifically and choose carefully per dataset (figures 2 and 3, supplementary figures 3 and 4).

The segmented right cheek unit has also been extracted as surface meshes (figure 4). The surface mesh data of the right cheek unit went through different stages of simplification process (Figure 4b-d) to produce good quality surface models with relatively small file size allowing quicker physical analysing (e.g. finite element analysis) and 3D printing. Mesh simplification results show that the number of vertices affects the presentation of the information and the detection of details (figure 3). When more triangles have been decimated, fewer details from data are detected (i.e. loss of information) resulting in a reduction in file size which allows easier sharing of the digital 3D model [20] or uploading the model to open data repository for educating the public [21] without giving out details that are still under investigation. By looking at the amount of details lost/preserved pointed by the arrows (figure 4), we recommend a mesh smoothing factor of 2 and 50% decimation based on our segmented cheek 3D model (figure 4d) as it halves the data size but still preserves most of the information. 3D models can then be used to generate 3D printouts [30] or 3D portable documents [31] to test a previous hypothesis or help with functional morphology investigations.

**Figure 4.**
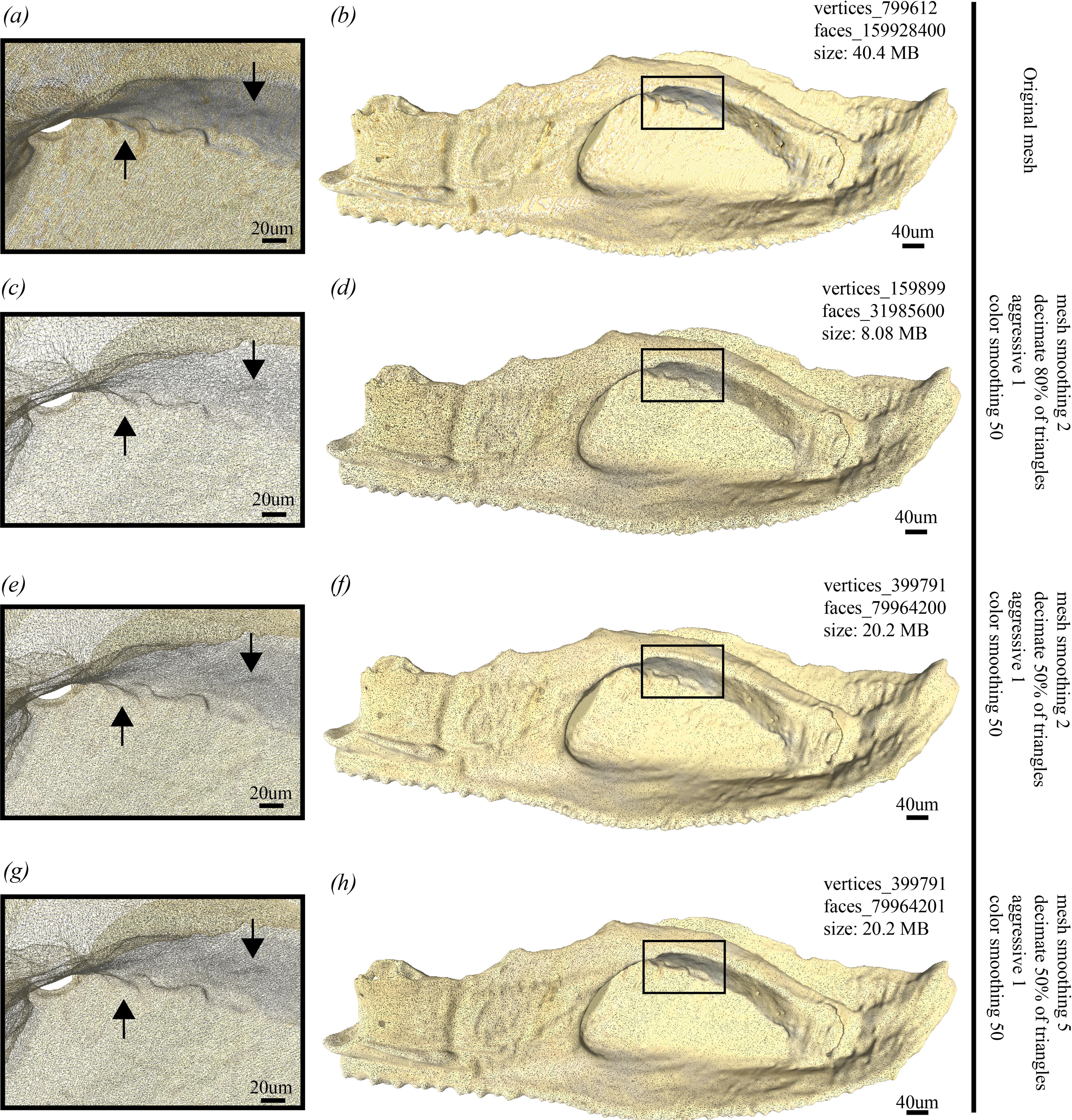
3D surface meshes of the extracted right cheek unit (Devonian placoderm fish, ANU V244) with attached perichondral-ossified cartilages, showing the differences of vertex count and file sizes after different mesh simplification applications. The Vertex count was calculated in MeshLab v1.3.4 [32].

## 4 Conclusions

Here we provide a new tool and workflow to segment volumetric data thus benefit the scientific community with more accurate and precise digital reconstruction, 3D modelling and 3D printing in *Drishti*. This procedure can be used as a framework to segment computed tomography and other forms of volumetric data and widely applicable in biological, medical and earth science researches. Our work will surely fuel further multiple cross-discipline collaborations between scientific visualization and many other fields.

## Data accessibility

3D surface mesh data of the segmented right cheek complex with different levels of simplification, and a movie shows the segmented internal canal networks of the toothplate are available from the fig share repository: https://figshare.com/s/244e5f407d331a39b402.

*Drishti* v2.6.6 is available from: https://github.com/nci/Drishti

## Authors’ Contributions

J.L. and Y.H. designed the study. Y.H., J.L., A.L. performed the research and drafted the manuscript. Y.H. and J.L. prepared the figures. A.L. developed *Drishti* v2.6.6. All authors revised the manuscript. Y.H. and A.L. contributed equally.

## Acknowledgements

We acknowledge Tim Senden and Gavin Young for their continuous support and help with proofreading. We thank the CT Lab at the Australian National University for CT Scanning.

