## Supplementary material for "A new tool for 3D segmentation of computed tomography data: *Drishti Paint* and its applications": SI text

**This supplementary Information includes:**

Supplementary Notes

Supplementary Table

Supplementary Figures

### Supplementary Notes

*Drishti*, as part of the service tools for scientists, has also been a part of this emerging new virtual science world (Supplementary Figure 1). From 2012 to the end of 2018, there are more than 14 fields of studies in physical sciences have been experienced with the program and benefited from the accessibility, transparency and ability to reproduction of a scientific analysis digitally through segmentation.

In this supplementary information, we will cover basic installation, user resources, basic background and supported data formats (Supplementary Table 1) for *Drishti*.

### *Drishti* v2.6.6

#### Installation

*Drishti* v2.6.6 is available to download from: https://github.com/nci/*Drishti* under "Releases".

When you click, it will lead to a version-specific webpage where the.zip file is located at the bottom of the page, click *Drishti* v2.6.6.zip to download *Drishti*.

Source code and detailed information about this new release are also available on Github. Please note that *Drishti* v2.6.6 currently can only run on the Windows operating system. However, users can compile and install *Drishti* v2.6.6 for CentOS/Ubuntu. Sample compilation script for *nix systems is provided to users with the source code.

Once the download is finished, unzip the .zip file. Go to "bins" folder, then you should be able to run *Drishti* v2.6.6. *Drishti* is designed as a portable application thus making sure you know where the portable directory stores on your computer.

#### MIT License

Copyright (c) 2011-2020 National Computational Infrastructure, Australia.

#### Other resources

Users can assess helpful videos through *Drishti* YouTube channel: https://www.youtube.com/channel/UCIomt3mEje4mGj_fwCNpxbQ

Tutorials and open datasets are available for practice: https://cloudstor.aarnet.edu.au/plus/s/ykqMmmikfXxHxKC?path=%2FTutorials

### *Drishti Import* v2.6.6

*Drishti Import* v2.6.6 converts the source data into the format (. pvl.nc) that Render (8 or 16 bits per voxel) and Paint (8 or 16 bits per voxel) read. *Drishti Import* v2.6.6 can read and convert 14 different file formats (see Figure 1 main text).

The main window (Supplementary Figure 2) is divided into two windows - the histogram window and the image window. The histogram window is used for the display of 1D histogram of the loaded volume (on the top). The image window displays the currently selected slice of volume.

***Drishti Paint v2.6.6***

*Drishti Paint* is designed for segmentation and data cropping of volumetric data. *Drishti Paint* v2.6.6 (Supplementary Figure 3) allows users to segment semi-automatically/manually and generate surface mesh for regions from the volume. To facilitate the segmentation process, *Drishti Paint* v2.6.6 provides 2 modes - Graph Cut and Curves. *Drishti Paint* v2.6.6 has been updated and now allows input of 16 bits full resolution volumetric data. More details of the segmented results of the internal canal networks of the selected region-of-interest on the right anterior upper toothplate of ANU V244 are showed in Supplementary Figure 4.

### Supplementary Table

| Data format | Name or supporting information |
| --- | --- |
| *.raw | ProRay raw triangle format;<br>a basic file format that saves the active image or stack as raw pixel data without a header |
| *. NIFTI | Neuroimagine Informatics Technology Initiative |
| TXM | Xradia X-ray transmission image data |
| *.jpg | Joint Photographic Experts Group |
| *. png | Portable Network Graphics |
| *.gif | Graphic Interchange Format |
| *.tiff/ Grayscale TIFF image | a tagged image file format |
| MetaImage | a special medical image format used in the insight segmentation and registration toolkit (ITK); |

| | |
|---|---|
| | a text-based tagged file format |
| Analyze 7.6 data format | Data format from a software named Analyze, which is developed by the Biomedical Imaging Resource at Mayo Clinic |
| *.nrrd/ NRRD | a library and file format designed to support scientific visualization and image processing involving N-dimensional raster data |
| *.txm/ QMUL Tom, TXM | a file format used by ZEISS Xradia 3D X-ray Microscopes |
| VGL | VG Studio Project file format |
| DICOM | Digital Imaging and Communications in Medicine<br>a standard medical imaging file format |

**Supplementary Table 1.** A summary of formats of volumetric data can be processed by *Drishti* v2.6.6.

### Supplementary Figures

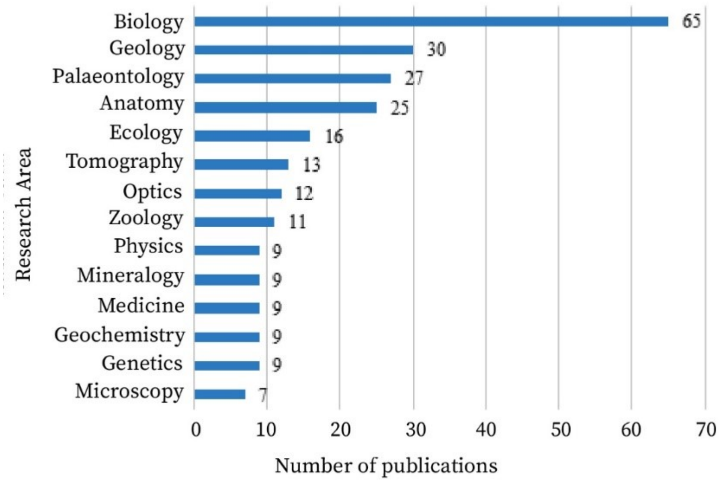


**Supplementary Figure 1.** Histogram of the number of publications used *Drishti* after Limaye 2012, publications categorized per research area/topic. Original data from Google Scholar and Microsoft Academic. Data analysed using Microsoft Excel 2018.

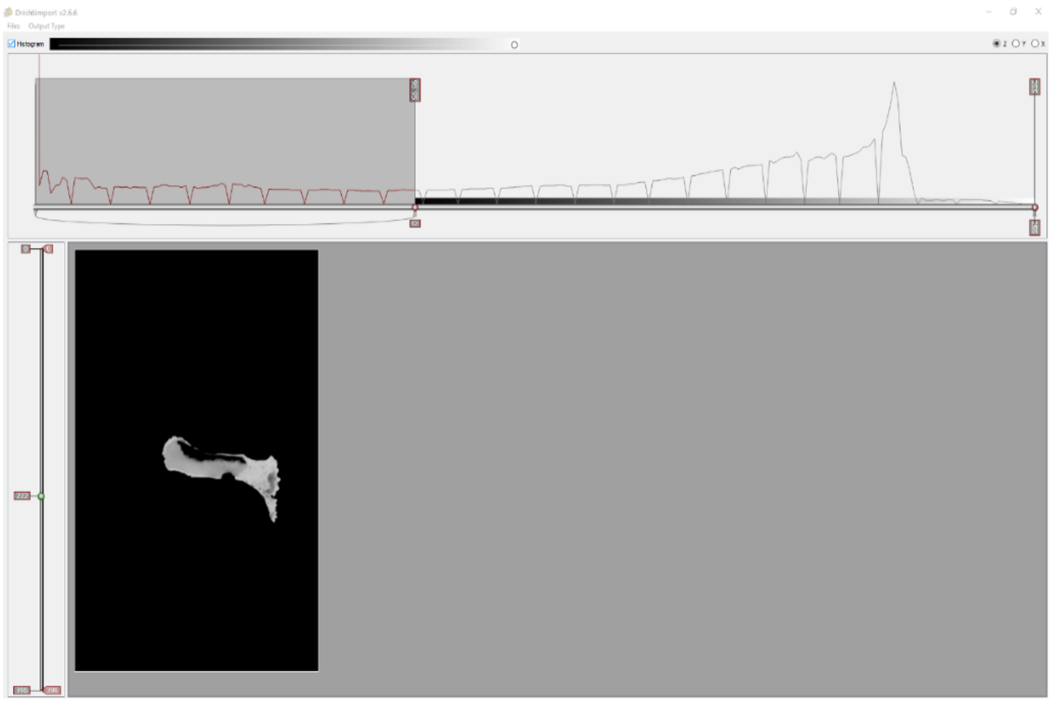

**Supplementary Figure 2.** The main window for *Drishti Import* v2.6.6.

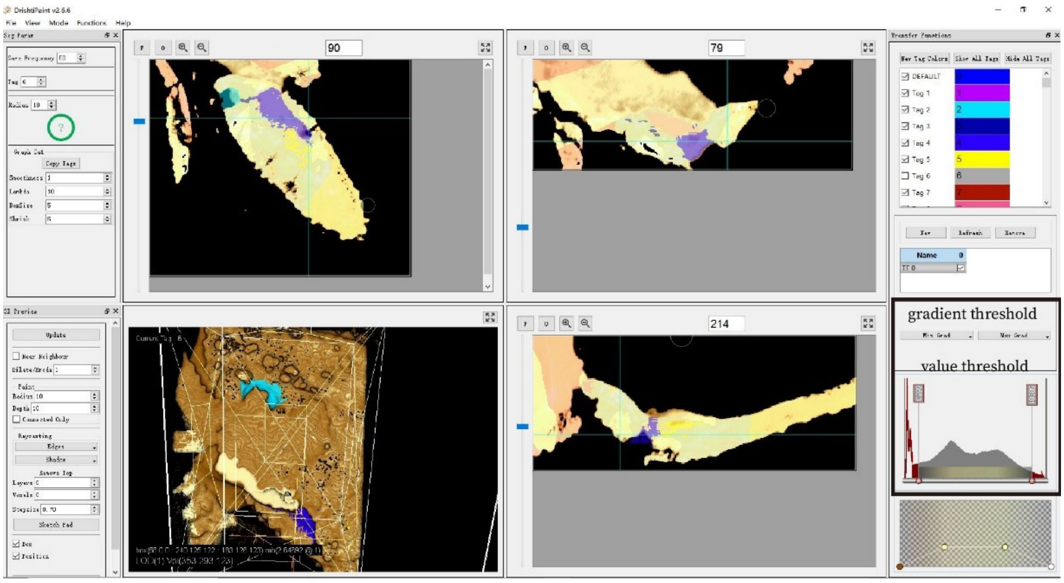


**Supplementary Figure 3.** The user interface of *Drishti Paint* v2.6.6 with newly developed value and gradient multi-thresholding for both Graph Cut and Curves modes, highlighted in the black box on the right side.

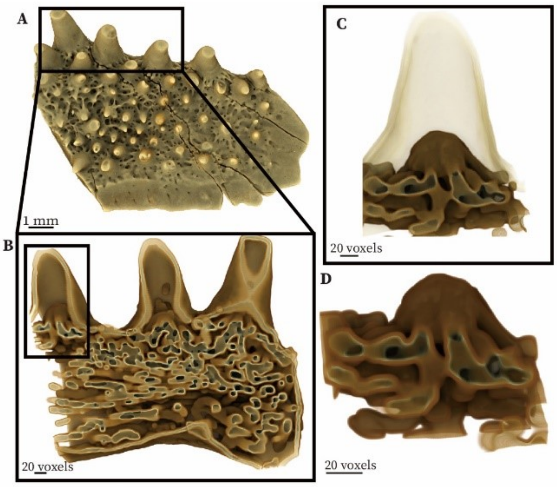


**Supplementary Figure 4.** Right anterior upper toothplate of a Devonian placoderm fish (ANU V244). A. Raw data loaded in *Drishti Import* v2.6.6.  B. Segmented results for the region-of-interest on the toothplate in *Drishti Render* v2.6.6. C. Single denticle with its outer surface and inner structure in *Drishti Render* v2.6.6. D. Inner structure of a single denticle in *Drishti Render* v2.6.6
